# Modeling and Minimization of dCas9-Induced Competition in CRISPRi-based Genetic Circuits

**DOI:** 10.1101/2025.11.05.686856

**Authors:** Dhruv D. Jatkar, M. Ali Al-Radhawi, Christopher A. Voigt, Eduardo D. Sontag

**Affiliations:** Department of Bioengineering, Northeastern University, Boston, MA, USA; Department of Biological Engineering, Massachusetts Institute of Technology, Cambridge, MA, USA; Department of Electrical and Computer Engineering, Northeastern University, Boston, MA, USA

## Abstract

Implementing logic functions in living cells is a fundamental area of interest among synthetic biologists. The goal of designing biochemical circuits in synthetic biology is to make modular and tractable systems that perform well with predictable behaviors. Developing formalisms towards the design of such systems has proven to be difficult with the diverse retroactive effects that appear with respect to the context of the cell. Repressor-based circuits have various applications in biosynthesis, therapeutics, and bioremediation. Particularly using CRISPRi, competition for components of the system (unbound dCas9) can affect the achievable dynamic range of repression. Moreover, the toxicity of dCas9 via non-specific binding inhibits high levels of expression and limits the performance of genetic circuits. In this work, we study the computation of Boolean functions through CRISPRi based circuits built out of NOT and NOR gates. We provide algebraic expressions that allow us to evaluate the steady-state behaviors of any realized circuit. Our mathematical analysis reveals that the effective non-cooperativity of any given gate is a major bottleneck for increasing the dynamic range of the outputs. Further, we find that under the condition of competition between promoters for dCas9, certain circuit architectures perform better than others depending on factors such as circuit depth, fan-in, and fan-out. We pose optimization problems to evaluate the effects engineerable parameter values to find regimes in which a given circuit performs best. This framework provides a mathematical template and computational library for evaluating the performance of repressor-based circuits with a focus on effective cooperativity.

## 1 Introduction

Since the discovery of the ability of genes to form circuits and perform logical operations [1], there has been an intense activity to build synthetic genetic circuits for various applications [2, 3, 4]. However, this field has faced significant obstacles due to the difficulty in characterizing the behavior of genetic logic gates, the limited number of orthogonal repressors, and many others [5, 6, 7].

Recent advances have mitigated some of these difficulties. In particular, the CRISPR-Cas9 system has been modified by deactivating Cas9’s endonuclease activity [8]. The resulting protein known as dCas9 maintains the ability to bind to a single guide RNA (sgRNA). The resulting complex binds to a specific area in the genome that matches the sgRNA. If the targeted area corresponds to a gene promoter, then the expression activity of the associated gene is turned off [9]. This new system has been nicknamed CRISPR interference (CRISPRi). The main advantage of CRISPRi is its programmability. This is since each dCas9-sgRNA complex binds the promoter that matches its sgRNA strand. Hence, this allows for the design of a very large number of mutually orthogonal repressors [10]. Some of the largest logic circuits to date have been built utilizing CRISPRi based on NOT and NOR gates [11].

However, despite its advantages, this technology suffers from significant bottlenecks that cause poor scalability. The first is dCas9’s toxicity which prevents it from being expressed at the levels required to support a large number of gates [12, 13, 14, 15, 16]. Second, its imperfect repression of the target promoters causes signal degradation in a cascade [10]. Approaches to mitigate this problem included the utilization of epigenetic mechanisms in eukaryotes like yeast to prevent expression leaks from repressed promoters [11], and reengineering dCas9 by fusing it to PhlF to increase its cooperativity and reduce its toxicity [14]. On the other hand, the aforementioned bottlenecks have motivated distributed-computing designs to scale-up the total number of gates [17, 18, 19].

Another set of approaches has focused on optimizing and regulating the performance of the circuit to minimize competition. Mathematical models have been proposed in [20, 14] which focused on parallel and series NOT gates. More recently, a closed-loop approach has been proposed for regulating the levels of dCas9 via a feedback controller [21]. Although it is a promising approach, implementing such regulators and predicting the behavior of a closed-loop high-dimensional and uncertain nonlinear system (such as a complicated genetic circuit) is not an easy task.

In this work we are motivated by the design of Boolean circuits and we focus on combinatorial logical circuits where the information will ideally flow in one direction. However, feedback effects are created by the competition for a limited dCas9 supply. Our mathematical analysis suggests that one of the major drawbacks of CRISPRi-based technology is its effective non-cooperativity. Although this issue has been discussed in the context of the difficulty of implementation of switches and oscillators [10],[22], its significance for the scalability of CRISPRi-based circuits is often overlooked. We suggest that the increase in circuit size in reengineered dCas9s [11],[14] can be interpreted in terms of the increase in effective cooperativity. Therefore, we suggest that the road to scalable CRISPRi runs through a more effectively cooperative dCas9.

## 2 Modeling a CRISPRi network

### 2.1 Modeling of the gates

We assume an arbitrary feedforward genetic circuit that consists of gates *G*_1_, …, *G*_*N*_ and external inputs *u*_1_, …, *u*_*m*_. Each gate *G*_*j*_ is a NOR gate with one or more inputs, and is *uniquely* associated with an sgRNA *s*_*j*_ and an output promoter *P*_*j*_ for *j* = 1, …, *N*. A NOR gate with a single input is called a NOT gate. We adopt the promoter-based formalism [23] where each gate maps promoter activities to promoter activities.

#### The NOT gate

Let *G*_*j*_ be a node in the CRISPRi network which has a single input *U*_*j*_. This input can be either an external input signal (i.e., *U*_*j*_ = *u*_*k*_ for some external input *u*_*k*_) or the output of another gate *G*_*i*_ (i.e., *U*_*j*_ = *P*_*i*_). The output of gate *G*_*j*_ is *P*_*j*_, which can drive other gates or serve as an external output. The protein dCas9 acts as a shared resource. A schematic diagram illustrating the gate is given in Figure 1(b).

**Figure 1:**
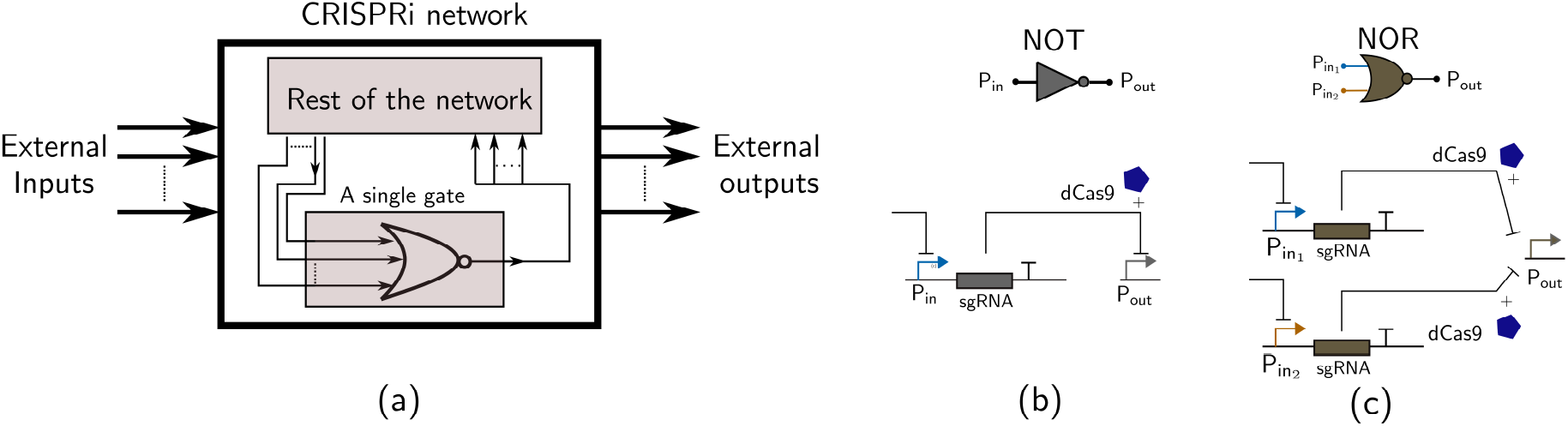
CRISPRi networks. (a) A general CRISPRi network. Each node in the network is a NOR/NOT gate. (b) A NOT gate. An active input promoter represses the output promoter. (c) A NOR gate with two inputs. The activity of either of the input promoters is sufficient to repress the output promoter. Note that the two inputs of the gates act on two different copies of the same sgRNA gene.

To model the system in Figure 1(a), we start by writing a reaction network model [24]. Let *s*_*j*_, *c*, and *C*_*j*_ denote the concentration of sgRNA, free dCas9, and sgRNA-dCas9 complex, respectively. Let *U*_*j*_ be the input promoter activity level. Then:

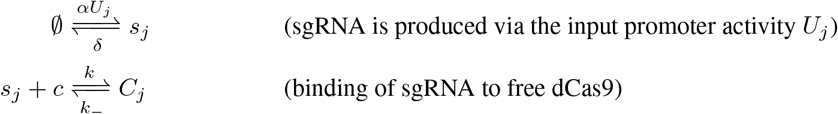

where *α, δ, k, k*_−_ are the kinetic parameters.

Assuming a Shea-Ackers cooperative binding model as in [25], we obtain the following model that characterizes the steady-state behavior of the gate:

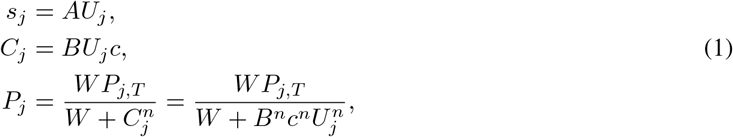

where 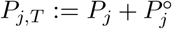 is the total amount of the *j*th promoter, 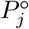 is the concentration of the bound promoter, *A*:= *α/δ, B* = *A/K, K*:= *k*_−_*/k, W* is a positive constant, and *n* is an effective cooperativity parameter.

The input to the gate can be proportional to the concentration of an external inducer, i.e., *U*_*j*_ = *u*_*k*_ for some external input *u*_*k*_, or equal to the output of another gate *G*_*i*_, i.e., *U*_*j*_ = *P*_*i*_.

The gate functions as a NOT gate for any *n* ≥ 1 for appropriately chosen *U*_*j*_. If *U*_*j*_ = 0, then *P*_*j*_ achieves the maximum possible value *P*_*j*_ = *P*_*j,T*_, and if *U*_*j*_ = ∞, then *P*_*j*_ = 0. In other words, the gate functions as a simple inverter.

Even if there are no dCas9 limitations, physical inputs cannot be infinite, and it is difficult to achieve zero inputs due to *promoter leakiness*. To model the latter phenomenon, when gate *G*_*j*_ receives input from gate *G*_*i*_, we can write:

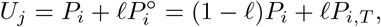

where *ℓ* ∈ [0, 1] is the leak percentage. The ideal case is *ℓ* = 0, while *ℓ* = 1 indicates a fully dysfunctional repressor.

#### A NOR gate

There are several ways to design a NOR gate. NOR gates can have an arbitrary number of inputs. For concreteness, here we discuss the case of two inputs. This can be done, for instance, by placing two promoters in tandem to control a single sgRNA gene. However, to reduce interference between the two promoters, a split design is preferred [10] as in Figure 1(c). Two copies of the same sgRNA gene are created, each controlled by a different input promoter. Therefore, the rate of production of sgRNA is proportional to the *sum* of the two input promoter activities. The only modification to (1) is:

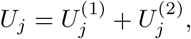

where 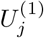 and 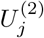 are the two input promoter activities to gate *G*_*j*_.

As in the previous case, the steady-state equations can be written similarly as:

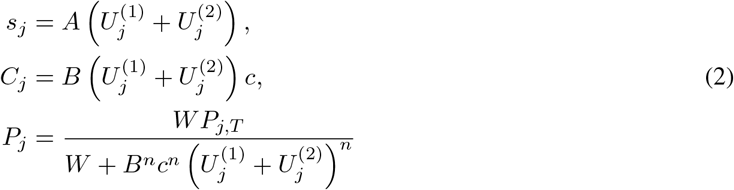

### 2.2 Modeling of competition

All the variables in Eq. (1)-(2) depend on the free dCas9 *c*. If it is assumed to be abundant, then *c* ≈ *c*_*T*_ where *c*_*T*_ is total amount of dCas9. However, if *c*_*T*_ is limited, then competition effects become apparent and the amount of free dCas9 allocated to each gate needs to be determined.

Once the free dCas9 is known, the level of dCas9 which is bound to each promoter can be found using (1),(2). It follows that the level of dCas9 allocated to each inhibited promoter follows a simple linear formula:

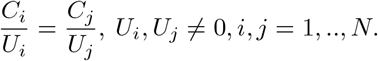

Writing the conservation law for dCas9, we get:

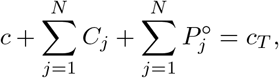

which can be written as (for given external inputs *u*_1_, .., *u*_*m*_):

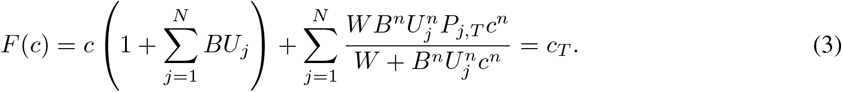

We call *F the conservation map*. Note that if *U*_*j*_ is equal to an output promoter, then some of the terms of the expansion above can be written as nested fractions. Therefore, the conservation map can be written in general as a *ratio of two polynomials*.

#### 2.2.1 Properties of the conservation map *F*

The following proposition states some of the basic properties of the function *F*:

##### Proposition 1.

Fix external inputs *u*_1_, .., *u*_*m*_. Let *F* be a function that takes the form (3). Then:

1. *F* is a rational polynomial of relative degree of 1, i.e., the degree of the numerator equals the degree of the numerator plus one.
2. *F* is onto, i.e., for any *c*_*T*_ *>* 0, let 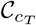 be the set of all positive solutions of (3); then 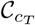 is nonempty.
3. For every given *c*_*T*_ *>* 0, let 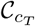 be the corresponding set of solutions of *F* (*c*) = *c*_*T*_; then 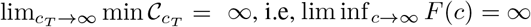.

*Proof*. The statement can be proved as follows:

1. Examining (3), it is sufficient to show that any term of the following form has a relative degree of zero:

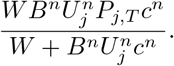

This can be proved by mathematical induction since inputs of each gate are rational polynomials of degree zero in terms of the outputs of the previous gates. We omit the details for brevity.
2. Since *F* has a relative degree 1, and the coefficient of the monomial of the highest degree is positive. Then *F* (∞) = ∞, i.e., lim_*c*→∞_ *F* (*c*) = ∞. Furthermore, *F* (0) = 0. By the intermediate value theorem, the equation *F* (*c*) = *c*_*T*_ has a positive solution for each *c*_*T*_ *>* 0.
3. We first show that 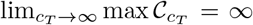. For the sake of contradiction, assume there exists *c*^*^ *>* 0 such that max 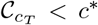 for all *c*_*T*_ *>* 0. Then this implies that *F* (*c*) *< c*^*^ for all *c* which contradicts the fact that *F* (∞) = ∞. In order to prove the statement, assume for the sake of contradiction that there exists *c*^*^ *>* 0 such that min 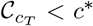 for all *c*_*T*_ *>* 0. Using the previous result, there must exist 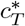 such that 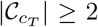 for all 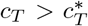. Using the Mean-value Theorem this implies that there exists an infinite number of points for which *dF* (*c*)*/dc* = 0. However, since *F*^*′*^ is another rational polynomial, it can not have an infinite number of zeros.

□

### 2.3 Quantifying and optimizing performance

#### 2.3.1 Quantifying performance

A Boolean genetic circuit represents binary zero and binary one by high and low promoter activity levels which are intrinsically analog. Therefore, we need a criterion to decide what ranges of value represent 1 and 0, and if there is sufficient “gap” between them so that they can be reliably distinguished.

Consider a given circuit with gates *G*_1_, .., *G*_*N*_, inputs *u*_1_, .., *u*_*n*_, and outputs *y*_1_, .., *y*_*m*_. Let 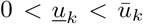 be the lowest and highest input levels for *k*th input *u*_*k*_, respectively. In other words, we have 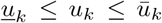 *k* = 1, .., *n* for all feasible choices of the inputs.

Let *f*_1_, .., *f*_*m*_: {0, 1} ^*n*^→ { 0, 1} be the desired Boolean functions that the circuit implements. For the *k*th input, the Boolean value 0 is represented by the inducer level 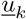 while the Boolean value 1 is represented by the inducer level 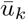. Therefore, for the purposes of evaluating the performance of the circuit, we can assume that each input is binary, i.e, the input vector 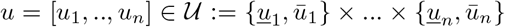 which we call the *input space*. Note that it is isomorphic to the Boolean input space {0, 1} ^*n*^. For ease of notation, we use the map *σ*: 𝒰 → {0, 1} ^*n*^ to record the transformation of physical concentration levels to logical values, and we denote its entries as *σ* = [*σ*_1_, .., *σ*_*n*_]^*T*^.

Let us consider the *p*th output *f*_*p*_(*σ*(*u*)). Note that the set of all possible inputs has cardinality 2^*n*^ and it can be divided into two mutually exclusive subsets. Hence, we define the sets <u>𝒰</u>_*p*_ and 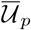 as follows:

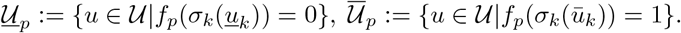

The sets 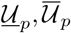 represent the combination of inputs that make the *p*th output low and high, respectively.

In order to measure the performance of the circuit with regards to the *p*th output, we need to distinguish between high and low levels of the output. Therefore, the following ratio must be as high as possible:

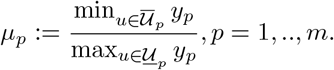

In order to evaluate the performance of the whole circuit, we write:

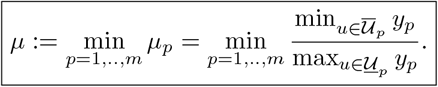

As used in the literature [14], we will require *µ* ≥ 10. In other words, we require that high outputs to be at least ten-fold larger than low outputs.

#### 2.3.2 Optimizing performance

Using our performance criteria and mathematical models, we can pose optimization problems that can be solved via *constrained nonlinear optimization* methods. We denote the optimization variables by *θ* which can include any parameters that can be modified. This can include promoter strength, gene dosage, RBS strength, target degradation, etc [26]. We consider two types of problems.

##### Minimizing total dCas9

Due to the well-documented toxicity of dCas9, it is desirable to minimize its total level while keeping the circuit functional.

Therefore, an optimization problem can take the form:

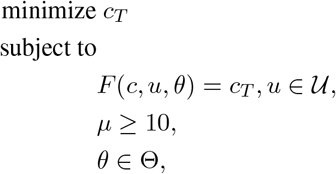

where Θ is the set of physical constraints that are imposed on the parameter set *θ*.

##### Maximizing performance

An alternative to problem of toxicity is to fix dCas9 to a non-toxic level, and then choose the optimization variables *θ* so as to maximize the performance index *µ*. Therefore, an optimization problem takes the following form for a given *c*_*T*_:

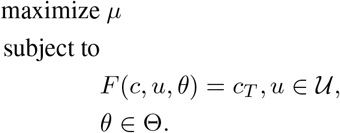

where Θ is the set of physical constraints that are imposed on the parameter set *θ*.

### 2.4 Uniqueness of steady states for two-input Boolean functions

The steady states of the presented genetic network satisfy the rational polynomial equation (3). Solving it amounts to solving a high-order polynomial equations in the cases of *n* = 1 and *n* = 2. Therefore, it is possible that such an equation admits multiple positive solutions. Next, we show that for the case *n* = 1 that network admits a unique positive solution for each given *c*_*T*_ *>* 0.

#### Proposition 2.

Consider a given a Boolean function with two-inputs. And let the corresponding circuit be a realization with a minimal number of gates. Let *n* = 1. Then the equations (3), (1), (2) admit unique positive solutions.

*Proof*. We prove this result by using deficiency-based algorithms [27] for each minimal circuit that realizes a two-input Boolean function.

For the case *n* = 2, the circuit realizations can admit multiple positive steady-states.

## 3 Examples

### 3.1 A cascade of two NOT gates

In order to illustrate the previous framework, let us consider a cascade of two NOT gates which should ideally function as a “repeater”, i.e., if the input is high, then output is high, and if the input is low the output is low. Therefore, we have *G*_1_, *G*_2_ where *U*_2_ = *P*_1_.

The input of the circuit is *u*, output of the circuit is *P*_2_ which can be written as:

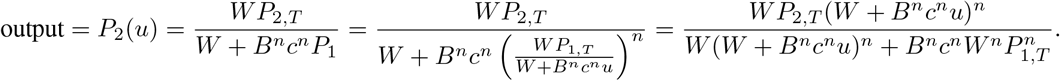

Let us examine first the “ideal” behavior of the circuit when *c* is abundant, zero promoter leak, and arbitrarily high inducer levels. We have:

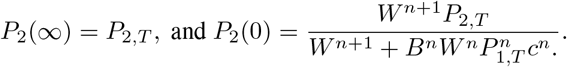

Therefore, the performance index *µ*_ideal_ is:

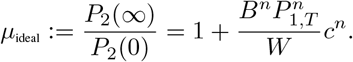

So even in idealistic conditions, a sufficiently high total dCas9 is needed to make *µ* ≥ 10.

However, in a resource constrained scenario, we need to solve for the free dCas9 using the conservation map *F*. Hence, we get:

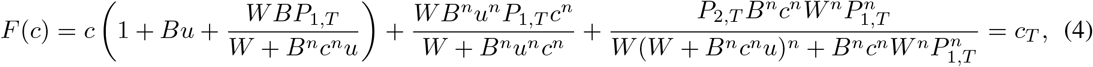

Solving for *c* amounts to solving a fourth order polynomial for the case *n* = 1, and a ninth order polynomial for *n* = 2. We use MATLAB’s command `roots` to carry this task numerically. For non-integer *n*, we use Netwon-Raphson’s method to find the solutions.

#### 3.1.1 The impact of cooperativity of the repressor

We first study how effective cooperativity affects performance. We use the following parameters: *B* = 0.03 *nM/s, W* = 17 *nM*, and *A* = 0.023 *nM/s* [14], and assume a 1% leak. Figure 2 compares the performance between *n* = 1 and *n* = 2. It can be noted that 33 times more dCas9 is required in the non-cooperative case to achieve the same performance as the cooperative counterpart. Since dCas9 is non-cooperative (*n* = 0.9 according to [14], this suggests strongly that the effect of competition can be mitigated by increasing the cooperativity of the repressors.

**Figure 2:**
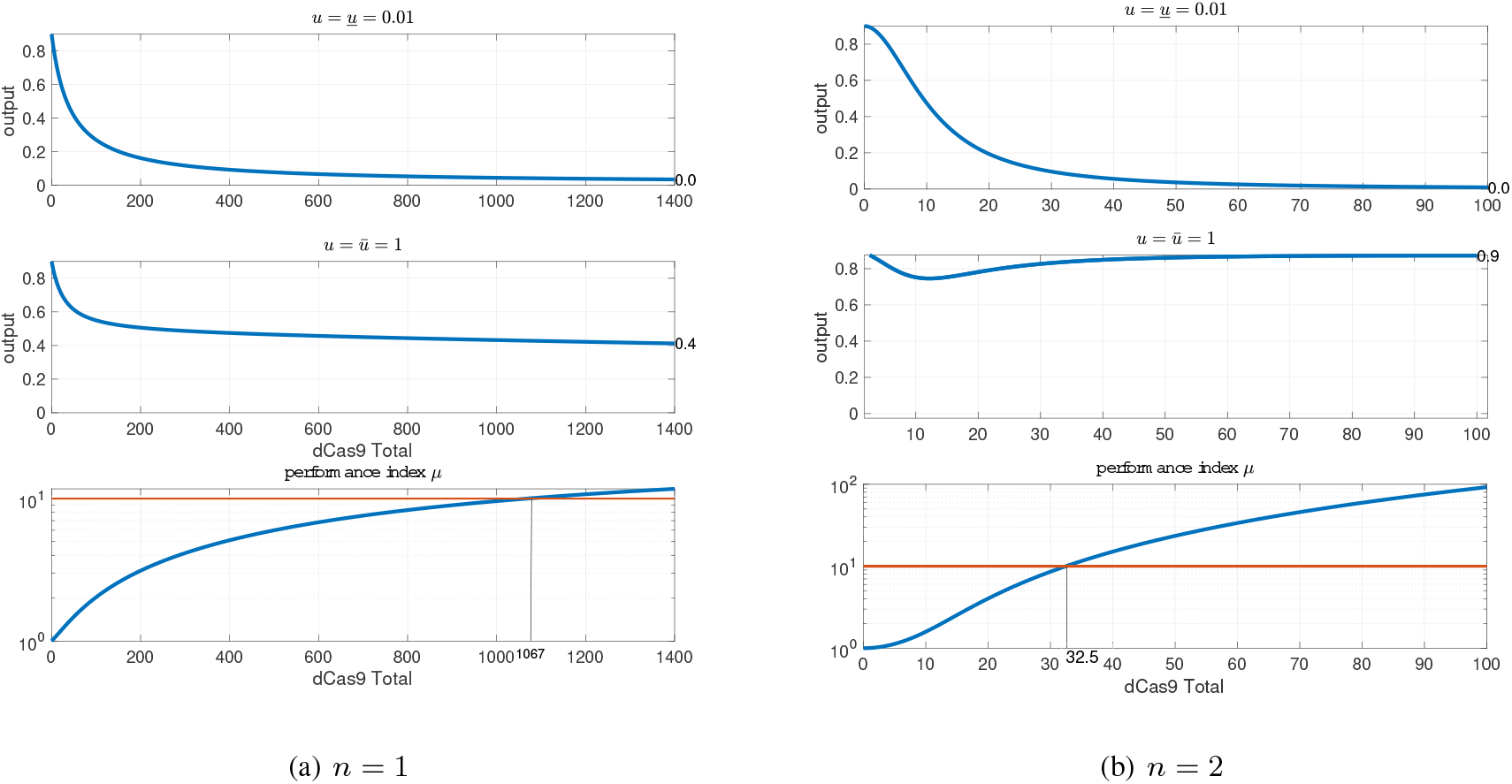
The performance of NOT-NOT network with different cooperativities.

#### 3.1.2 The impact of promoter copy-number

The optimization framework can be utilized to maximize the performance over manipulable variables. For instance, consider that the gene dosage or the gene copy number *P*_1,*T*_ for the NOT-NOT circuit as an optimization variable.

Then, we solve the following optimization problem to find *P*_1,*T*_, *c*_0_, *c*_1_ for a given *c*_*T*_, *ū, <u>u</u>*:

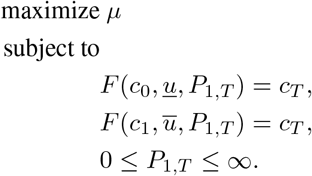

In the previous subsections, we have shown that in order to achieve performance level *µ* = 10 we need a minimum of *c*_*T*_ = 32.5 with *n* = 2 and the total promoter copy number for the first gene assigned to *P*_1,*T*_ = 1.

Solving the optimization problem numerically yields that the optimal promoter copy number is *P*_1,*T*_ = 1.99, and the optimal performance index is 15.266 which is a 50% improvement.

#### 3.1.3 Impact of leak

An sgRNA-gate functions by repressing the target promoter. However, such promoters can remain partially active under repression. Such phenomenon is known as *promoter leakiness*, and it is one of the major hurdles against the scalability of genetic circuits [26].

In the context of CRISPRi, increasing the total amount of dCas9 without bounds means that a leaky promoter gets a higher chance of being expressed. Hence, the performance of the circuit begins to deteriorate for higher *c*_*T*_. Therefore, there exists an optimal amount of *c*_*T*_ that depends on the leakiness level.

For instance, consider the NOT-NOT circuit with *<u>u</u>* = 0.05, 5% leak, and *n* = 2. We plot the performance index versus the input level and total dCas9 in Figure 3. It shows that a total amount of dCas9 that is higher than ∼100 requires higher input levels to maintain the performance.

**Figure 3:**
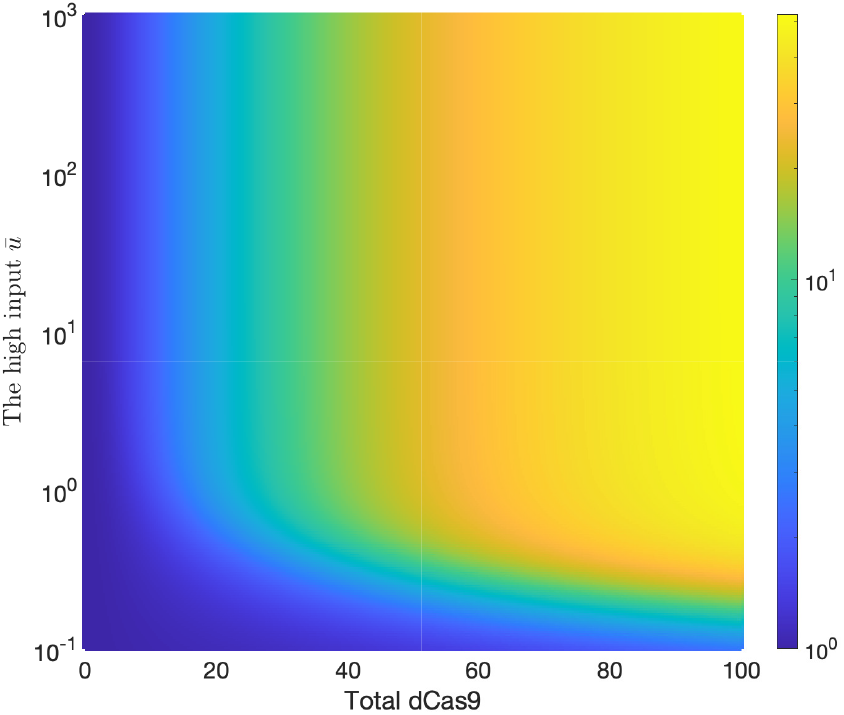
The performance of the circuit versus the input level and total dCas9 with high leak. The circle denotes the optimal operating point.

## 4 Impact of Circuit Architecture on Performance

While genetic circuits can be designed with multiple architectures that implement the same Boolean function, these architectures do not perform equally under various context effects, including resource competition. The topology of a circuit (how gates are connected and organized) fundamentally affects its behavior when dCas9 is limited. Understanding these architectural effects is crucial for rational circuit design.

We compare two different designs of the XOR gate. Each design consists of five gates and is depicted in Figure 4. We assume *<u>u</u>* = 0.05, 5% leak, and *n* = 1.6. Throughout all the simulations in this paper, we use the following parameters *B* = 0.03 *nM/s, W* = 17 *nM, A* = 0.023 *nM/s* [14]. Figure 4 shows the results for both designs. It can be seen that the first design is significantly more robust against leakiness compared to the second design. In the following sections, we consider three structural properties of genetic circuits. We define the depth of a circuit as maximum number of gates that an input can be processed through over all paths, to the output. We define the fan-in of a gate *G* as the number of upstream gates whose outputs are an inputs to *G*, and the average fan-in of the circuit as the sums of the fan-ins over all gates divided by the number of gates. We define the fan-out of a gate *G* as the number of downstream gates where the output of *G* acts as an input, and the average fan-out of the circuit as the sums of the fan-outs over all gates divided by the number of gates.

**Figure 4:**
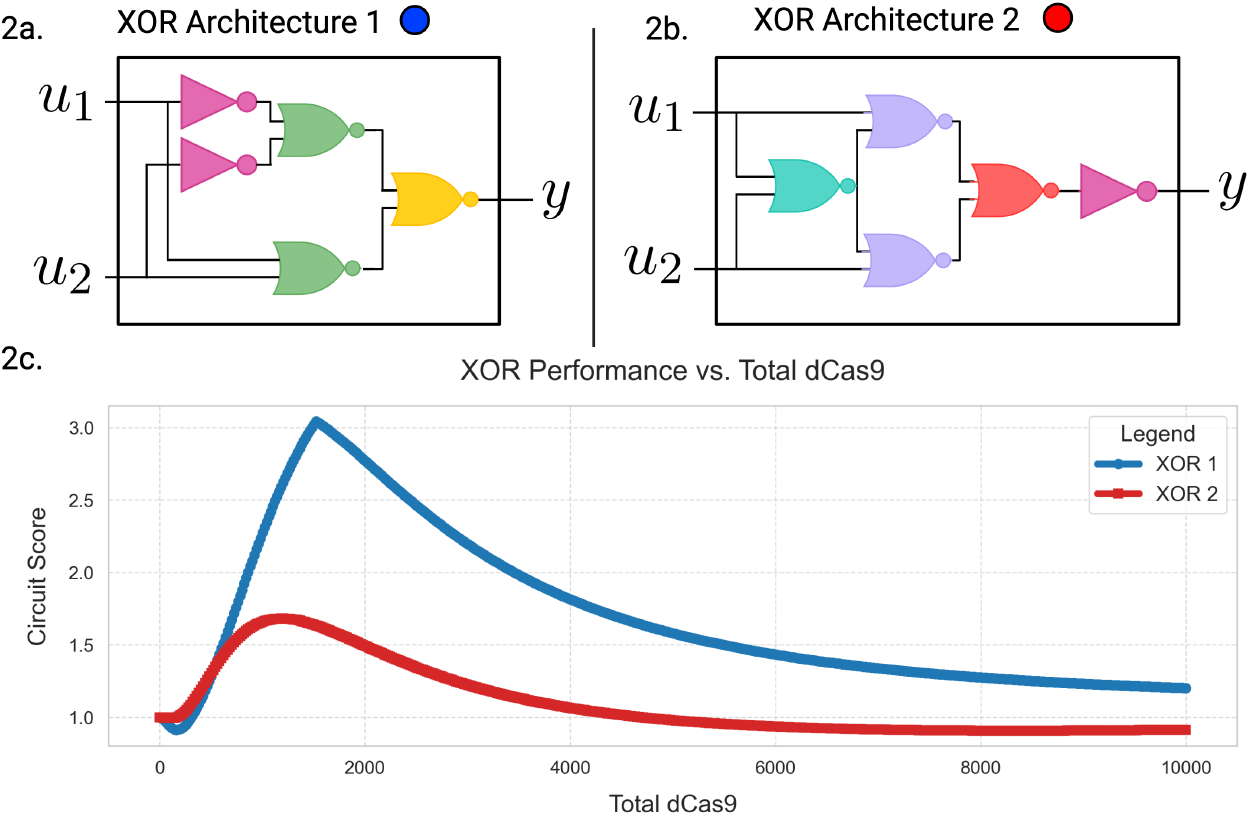
Different XOR. Architectures The XOR function has several different architectures. Comparing two architectures (a) and (b), we find that they perform differently when comparing circuit score to total dCas9 in (c).

### 4.1 Depth–induced signal degradation in CRISPRi cascades

Depth is the first architectural feature we decide to interrogate, and the one most studied in experimental literature. In our setting, every added layer introduces two compounding pressures. Firstly, a partially repressed promoter becomes the input to the next gate, causing the leak to propagate downstream. And second, all layers draw from the same finite dCas9 pool, so the free complex *c* allocated by the conservation law (3) decreases as more sgRNA species compete for it. Even if individual gates are well characterized in isolation ((1)-(2)), the global coupling through *c* means that the steady state of a deep cascade cannot be inferred by composing single-gate transfer curves alone. To isolate the effect of depth from other architectural variables, we analyzed a cascade of *d* ∈ {1, …, 7} NOT gates that share a single dCas9 pool. For each depth *d* and each total dCas9 level *c*_*T*_, we solved the conservation equation (3) at the two input settings that define the *ON* and *OFF* logical cases (for odd *d* the ON state uses *u* = *<u>u</u>*, for even *d* it uses *u* = *ū*), and we computed the circuit score *µ*. The resulting depth curves are shown in Fig. 5.

**Figure 5:**
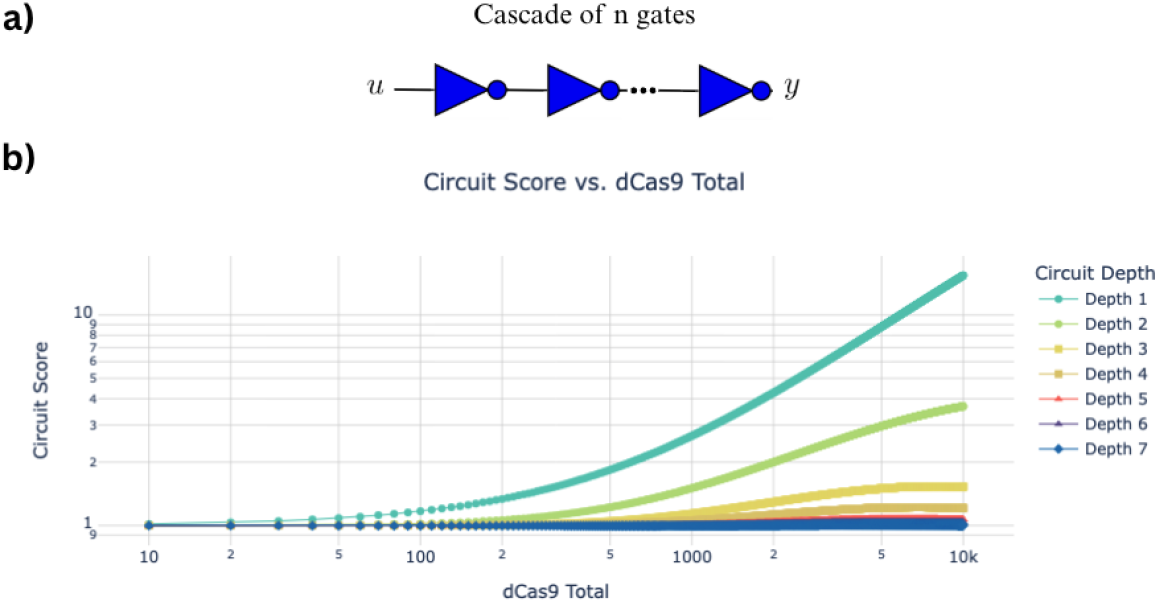
Depth-induced loss of circuit score in CRISPRi cascades under dCas9 competition. **(a)** Schematic of a cascade of *n* NOT gates sharing one dCas9 pool; input *u* drives the first gate and the output *y* is read at the last gate. **(b)** Circuit score *µ* (ratio of the steady-state output in the intended ON state to the output in the intended OFF state) versus total dCas9 *c*_*T*_ for cascades of depth 1–7 (log–log axes). ON/OFF are chosen by parity: for odd depth the ON case uses *u* = *<u>u</u>* = 0.01 and the OFF case uses *u* = *ū* = 1.0; for even depth the assignments are reversed. At any fixed *c*_*T*_, *µ* decreases monotonically with depth—deeper cascades suffer stronger signal degradation—and increasing *c*_*T*_ only partially recovers contrast over *c*_*T*_ ∈ [10, 10^4^]. Curves are computed from the promoter-based CRISPRi model with dCas9 conservation solved by Newton–Raphson at each *c*_*T*_. Simulations were computed at cooperativity *n* = 1 (non-cooperative repression); leak *ℓ* = 0.15; promoter total *P*_*T*_ = DT = 1; *W* = *w*_−_*/w* = 17 with *w* = 1, *w*_−_ = 17; *K* = *k*_−_*/k* = 1.08 with *k* = 1, *k*_−_ = 0.03*/δ* = 1.08; *B* = 0.03; *δ* = 100*/*3600, *γ* = 1*/*3600; scaling factor *fa* = 1; input levels *<u>u</u>* = 0.01, *ū* = 1.0; *c*_*T*_ swept linearly over 1000 points from 10 to 10^4^.

At any fixed *c*_*T*_, the circuit score *µ* decreases monotonically with depth. Raising *c*_*T*_ improves *µ* for each depth, but with diminishing returns: beyond a moderate *c*_*T*_ the curves flatten, and additional dCas9 yields only marginal gains. This saturation is a direct consequence of the conservation map *F* in (3): each additional layer introduces a new sgRNA pool that sequesters complex and creates more bound promoter sites, both of which depress the free *c* available to all other layers. In parallel, leak (*ℓ >* 0) is effectively amplified by depth because small residual outputs at one stage become inputs to the next. When *n* = 1, these two mechanisms combine to diminish *µ* as *d* grows.

However, depth is only one lever. In practice, designers often exchange depth for higher fan-in (using NOR gates that sum multiple promoter activities) or accept some fan-out to reuse intermediate signals. Both choices reshape the demand on dCas9: higher fan-in can strengthen repression in the intended OFF states (potentially boosting *µ*), while higher fan-out spreads a single promoter’s output across more consumers (typically hurting *µ*). The next section quantifies these trade-offs by enumerating architectures for two-input Boolean functions and aggregating circuit scores as a function of depth, fan-in, and fan-out.

### 4.2 Interplay between fan-in, fan-out, and depth

In the previous sections, we described specific ways in which a circuit can gain or lose performance with respect to circuit architecture. However, this does not always mean that if a circuit has a larger depth than another circuit, it will also have a lower circuit score. In fact, it is often the case that we observe trade-offs between depth, fan-in, and fan-out with certain structural features dominating in contribution to circuit score. To observe these effects we enumerate all 2-input 1-output Boolean functions. For each of these functions, we enumerate all satisfying architectures and compute the maximum circuit score over a realistic range of total dCas9. We aggregate the scores of each circuit, tracking circuit depth, fan-in, and fan-out to observe trends. In this case, we fix the number of gates in each circuit to 5.

In Figure 6, we find that in order to have circuits with a larger average fan-in, we must trade depth. This shows that while we may lose performance through an increased depth, fan-in is a dominating contributor to increasing performance.

**Figure 6:**
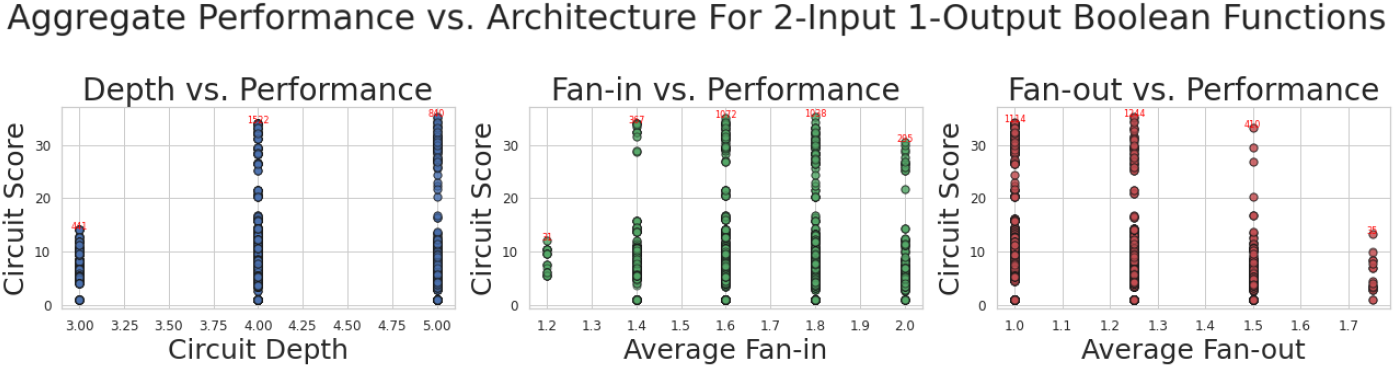
Aggregating Circuit Scores to Observe Trends. By exhaustively enumerating all 5-gate circuits for all 2-input 1-output Boolean functions, we are able to observe trends in circuit score with respect to structural features. We find that generally, higher fan-in increases circuit score while higher fan-out decreases circuit score. Additionally, the loss of performance through a deeper circuit is overshadowed by the larger average fan-in one can attain by increasing depth.

### 4.3 The competition model reproduces effects on circuit depth

Sections 4.1-4.2 established two modeling claims: (i) depth erodes contrast in CRISPRi cascades under limited dCas9, and (ii) architectural choices that increase fan-in (and restrain fan-out) can partially recover performance by reshaping how the network draws from the shared dCas9 pool. A natural next question is whether the same resource competition mechanism encoded by the conservation map *F* in (3) and the gate equations ((1), (2))is sufficient to explain experimentally measured depth effects without re-tuning parameters gate by gate.

We performed a single global calibration against the depth series reported for NOT-cascades in [14], fitting our competition model across depths *d* ∈ {1, 2, 3, 4} simultaneously (Fig. 7). For every input setting in each depth, the free dCas9 concentration *c* was obtained by solving the conservation equation (3); the resulting steady states were compared to published experimental data [14] in log space. Crucially, parameters were shared across depths, enforcing that the only way to reconcile all curves is through the same mass-balance constraint that couples gates via *c*. Our fits demonstrate that one set of biophysical parameters, together with explicit dCas9 conservation, explains the entire depth series. Practically, it means a laboratory calibration performed once (on a small panel of circuits) can be reused to predict the performance of new architectures without re-fitting every gate. Practically, we treat the conservation law (3) as a design guide. In our simulations, three levers generally help at fixed *c*_*T*_: increasing effective cooperativity *n*, reducing leak *ℓ*, and limiting fan-out (which lowers ∑ _*j*_ *BU*_*j*_). These changes tend to raise *µ*, although the size of the improvement depends on the circuit and parameter choices.

**Figure 7:**
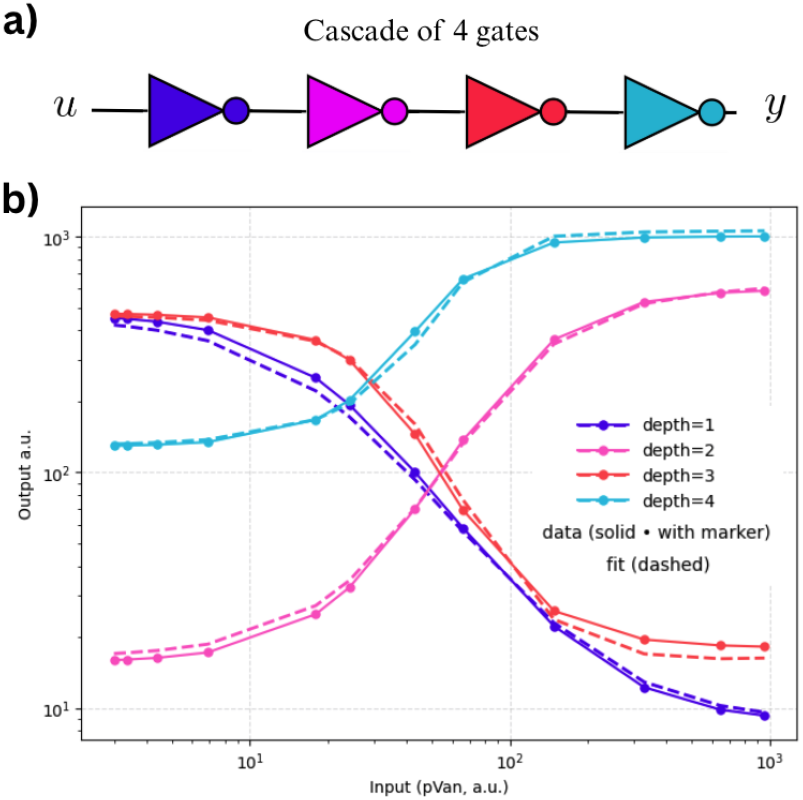
Depth–dependent response of a NOT-cascade; global competition-model re-fit of published data. **(a)** A four-stage NOT cascade from input *u* to output *y*. **(b)** Solid curves (with markers): cascade measurements reproduced from Fig. 2D of [14]. Dashed curves: a *single* global fit of the dCas9 competition model that enforces mass balance for free dCas9 across the entire cascade; the same parameter set predicts all depths *d* ∈ {1, 2, 3, 4}. Calibrating the competition model once (shared parameters across all depths) yields high-fidelity, depth-aware predictions of steady-state response curves, improving upon gate-by-gate compositions that ignore resource sharing. Together, these results show that an explicit resource-competition model predicts the entire family of depth-dependent responses with a single parameter set.

## 5 Discussion and directions

CRISPRi technology has the advantage in its programmability. A single dCas9 protein can carry a very large number of orthogonal sgRNA molecules each encoding the output of a single gate. However, the major obstacle with respect to scalability is the excessive demand for dCas9 in CRISPRi circuits. Such demand cannot be sustained in large circuits without compromising the host cell growth rate.

In this report, we present the results of our undergoing research that aims at building a computational framework to improve our understanding of resource competition in CRISPRi circuits. Our findings suggest that high cooperativity helps in reducing the demand for dCas9, and hence in building more scalable circuits.

In terms of future direction, we plan to develop this into a computational package, and do more detailed modeling of re-engineered dCas9 variants. In addition, we will be writing a follow-up report in which we mathematically prove the existence of the trends observed in Section 4.

## Acknowledgements

This work was partially funded by the Air Force Office of Scientific Research under award FA9550-22-1-0316 (DDJ, MA, EDS) and the National Science Foundation SemiSynBio program under award CCF-1849588 (CAV, EDS).

## A Enumerating architectures

### A.1 All architectures for every 5-gate 2-input Boolean function

In order to view the trends described previously between depth, fan-in, and fan-out, we perform simulations for all Boolean functions with *k* gates, 2-inputs, and 1 output for *k* = 3, 4, 5, 6, 7. For each function, we construct a truth-table for reference in a exhaustive search over every digraph representing *k* gates. If the corresponding circuit is valid, we compute the circuit-score over a range of *c*_*T*_ s.

Observe that from the simulations below, there exists a trade-off between circuit depth and average fan-in where the most optimal architecture often is a deeper circuit which allows for more NOR2 gates to have multiple inputs allowing for increased average fan-in.

**Figure 8:**
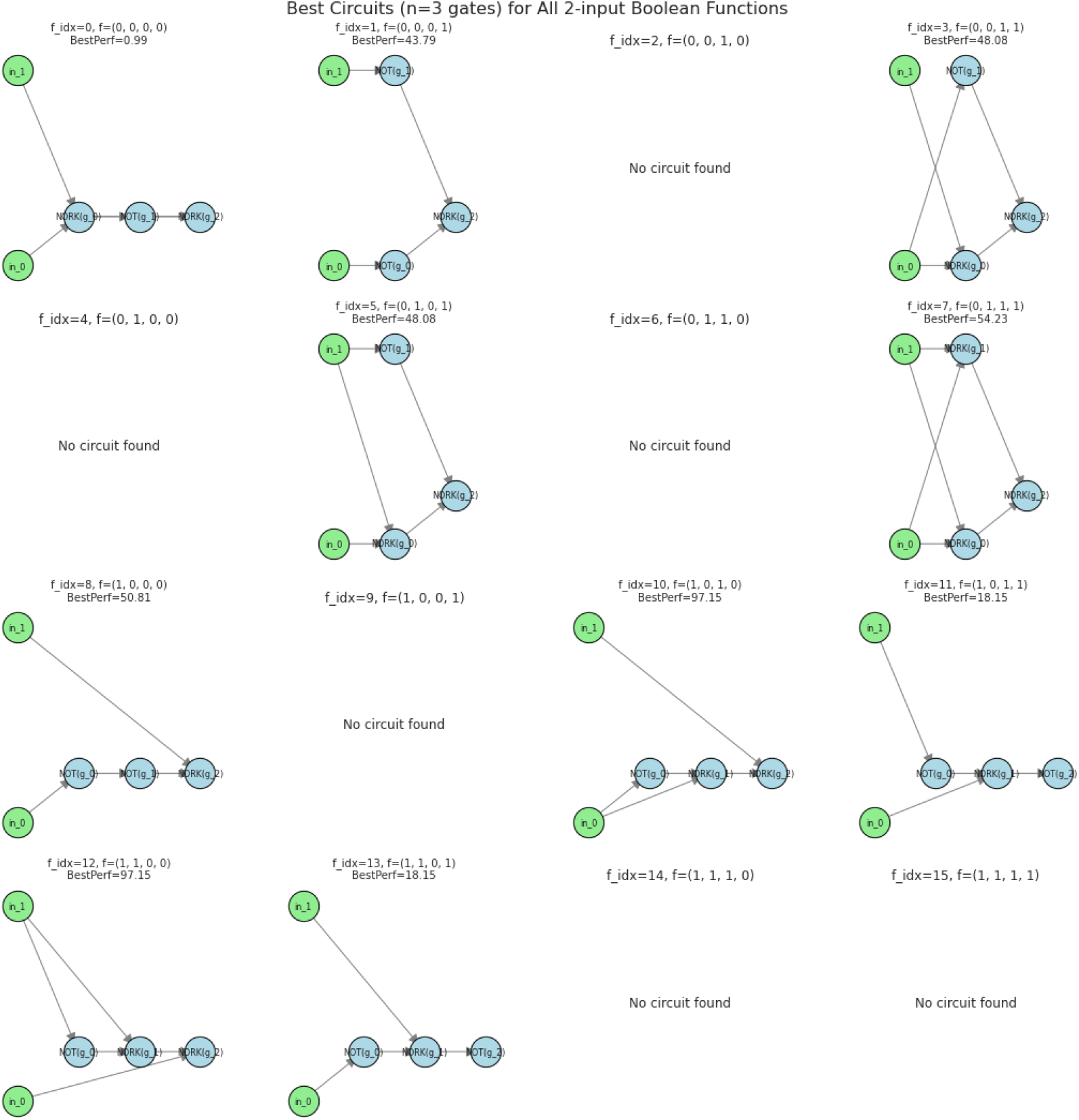
A. Optimal circuit architectures and *µ*s for all 2-input Boolean functions using *k* = 3 gates

**Figure 9:**
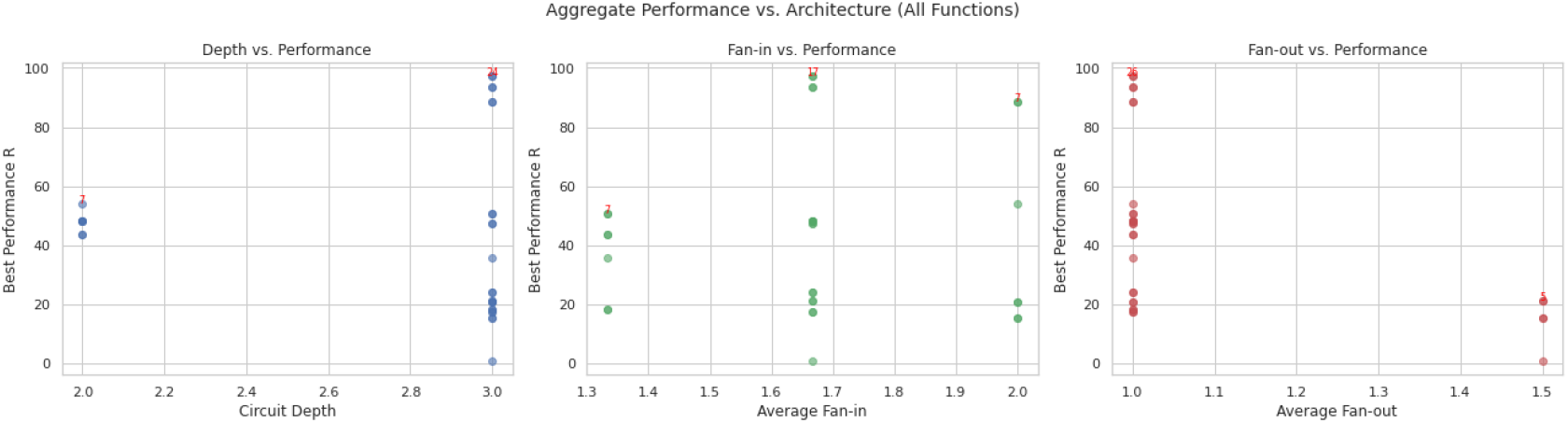
B. Aggregate plots comparing *µ* to circuit depth, average fan-in, and average fan-out for all 2-input Boolean functions using *k* = 3 gates

**Figure 10:** Over all 2-input Boolean functions using *k* = 3 gates, we observe that performance generally increases as average fan-in increases and average fan-out decreases. A higher depth allows for a higher average fan-in resulting in larger *µ* with a deeper circuit.

**Figure 11:**
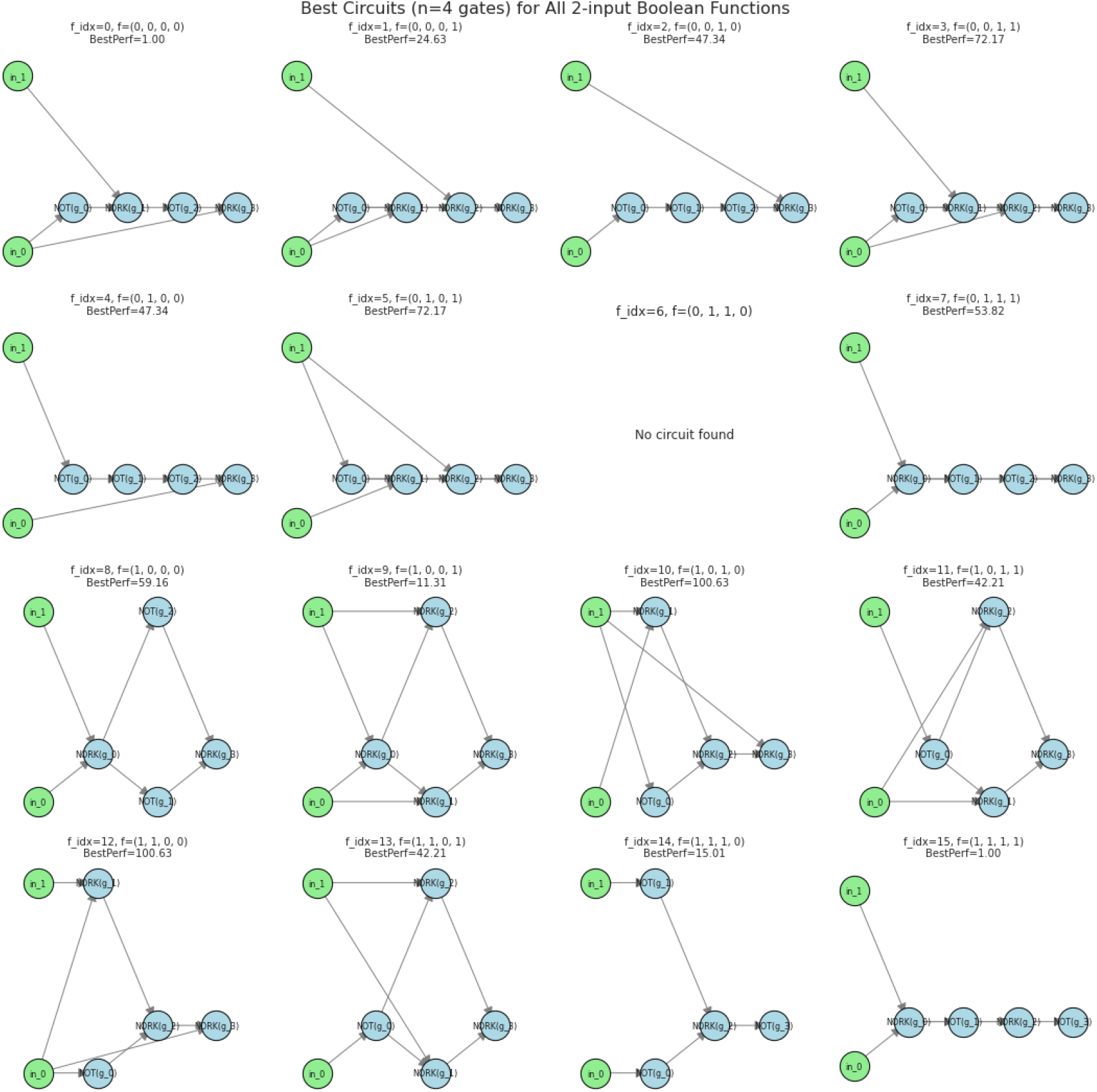
A. Optimal circuit architectures and *µ*s for all 2-input Boolean functions using *k* = 4 gates

**Figure 12:**
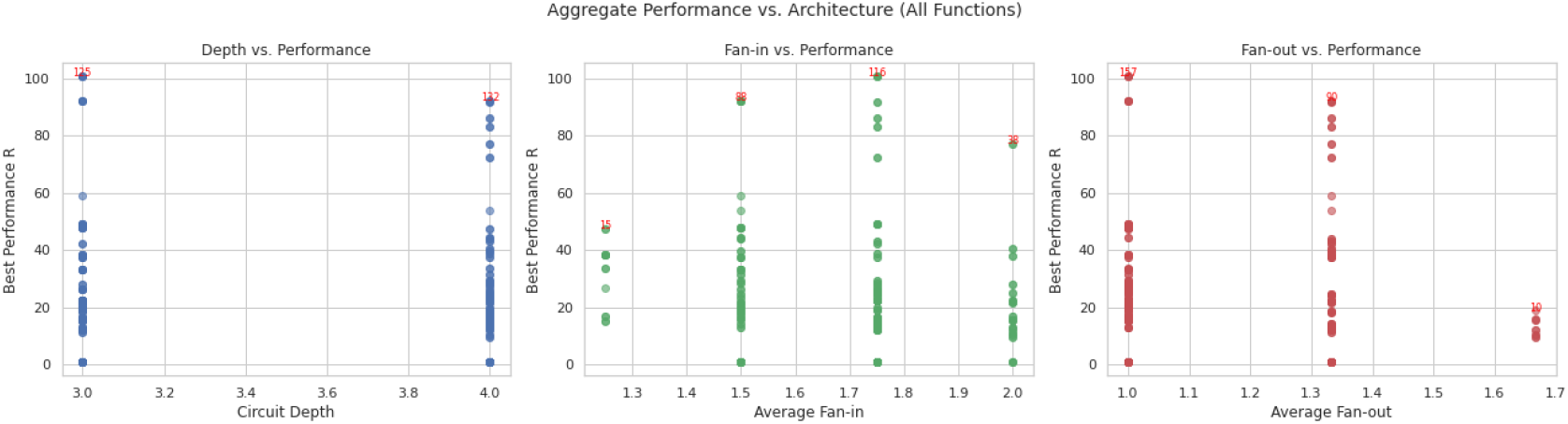
B. Aggregate plots comparing *µ* to circuit depth, average fan-in, and average fan-out for all 2-input Boolean functions using *k* = 4 gates

**Figure 13:** Over all 2-input Boolean functions using *k* = 4 gates, we observe that performance generally increases as average fan-in increases and average fan-out decreases. A higher depth allows for a higher average fan-in resulting in larger *µ* with a deeper circuit.

**Figure 14:**
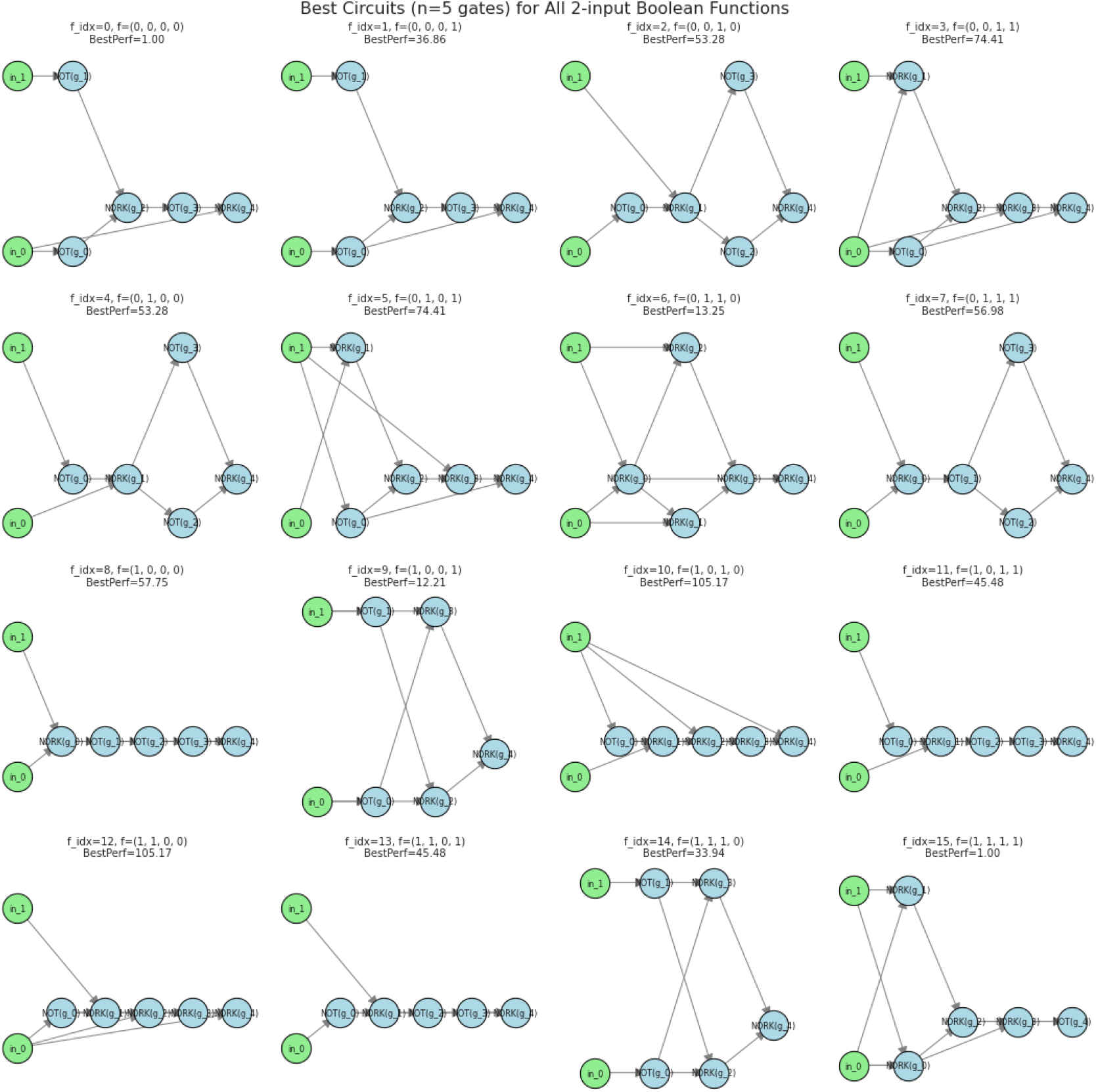
A. Optimal circuit architectures and *µ*s for all 2-input Boolean functions using *k* = 5 gates

**Figure 15:**
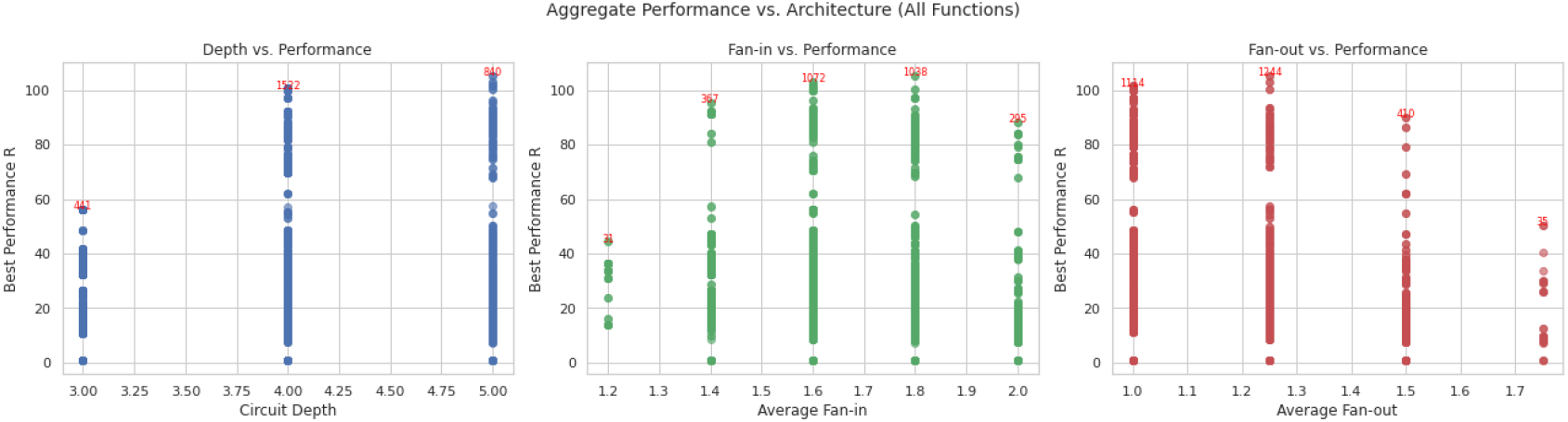
B. Aggregate plots comparing *µ* to circuit depth, average fan-in, and average fan-out for all 2-input Boolean functions using *k* = 5 gates

**Figure 16:** Over all 2-input Boolean functions using *k* = 5 gates, we observe that performance generally increases as average fan-in increases and average fan-out decreases. A higher depth allows for a higher average fan-in resulting in larger *µ* with a deeper circuit.

**Figure 17:**
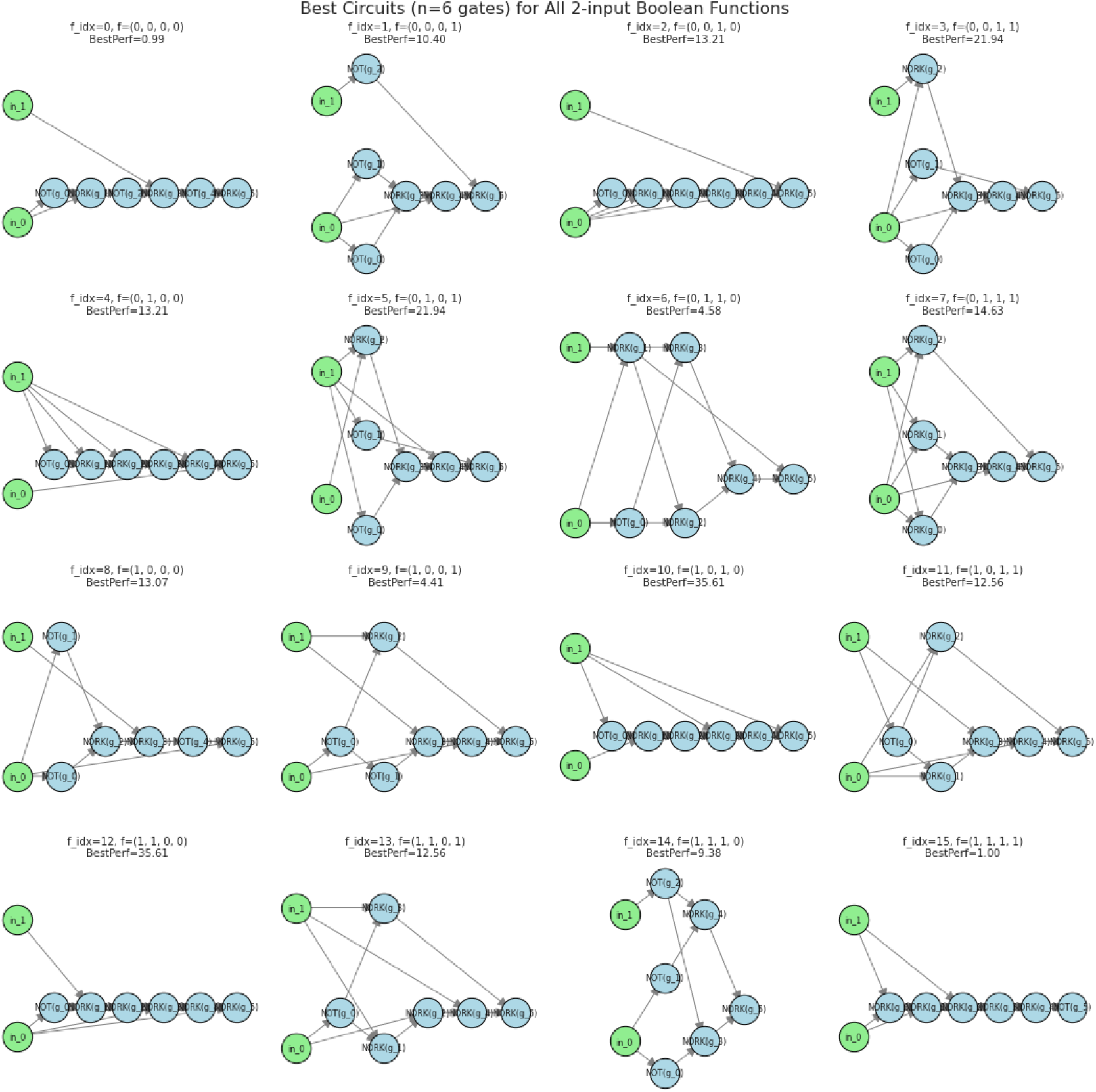
A. Optimal circuit architectures and *µ*s for all 2-input Boolean functions using *k* = 6 gates

**Figure 18:**
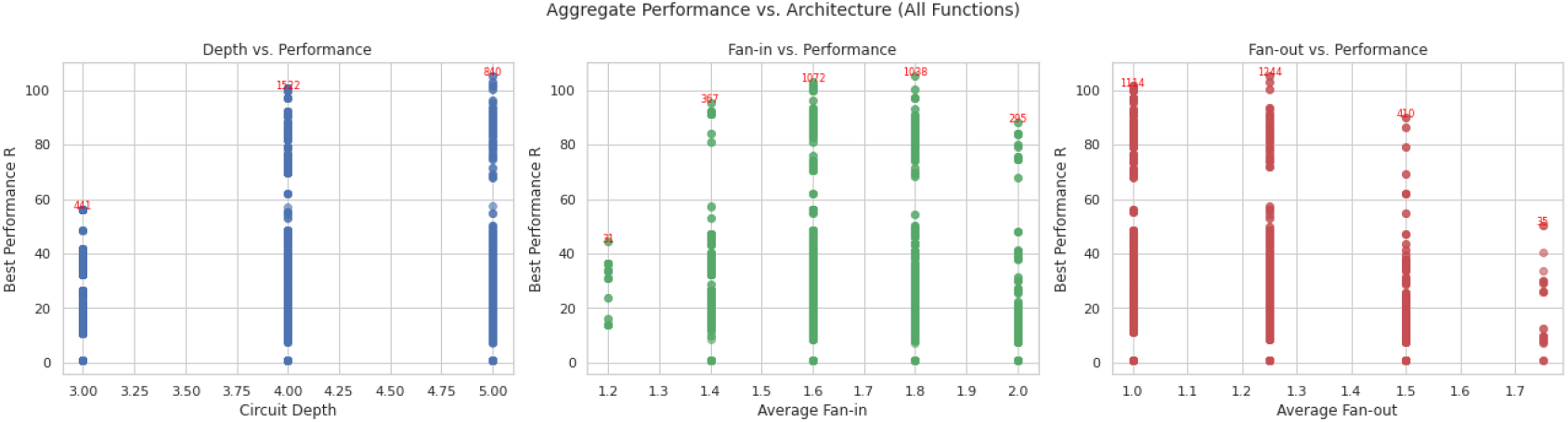
B. Aggregate plots comparing *µ* to circuit depth, average fan-in, and average fan-out for all 2-input Boolean functions using *k* = 6 gates

**Figure 19:** Over all 2-input Boolean functions using *k* = 6 gates, we observe that performance generally increases as average fan-in increases and average fan-out decreases. A higher depth allows for a higher average fan-in resulting in larger *µ* with a deeper circuit.

**Figure 20:**
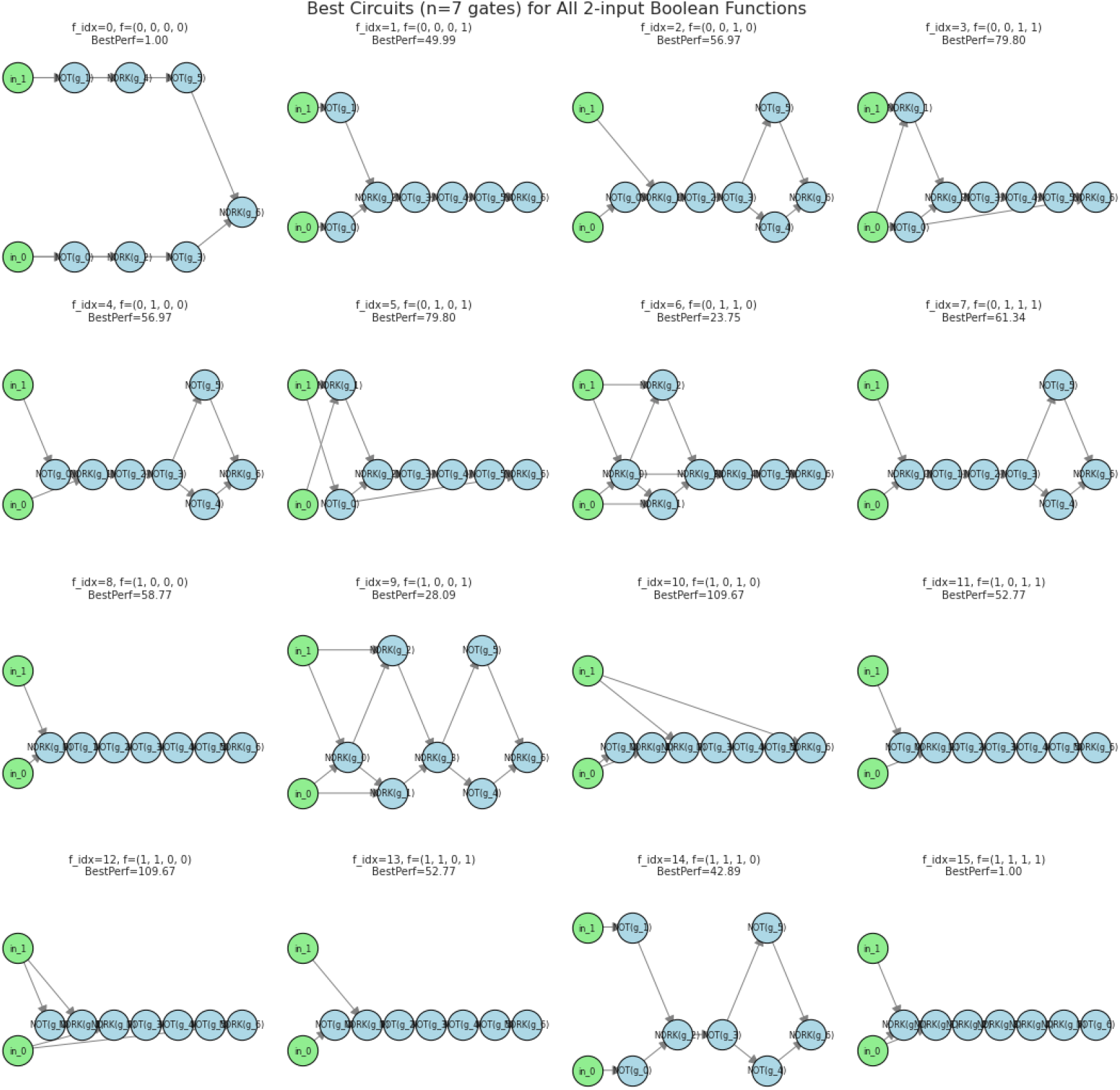
A. Optimal circuit architectures and *µ*s for all 2-input Boolean functions using *k* = 7 gates

**Figure 21:**
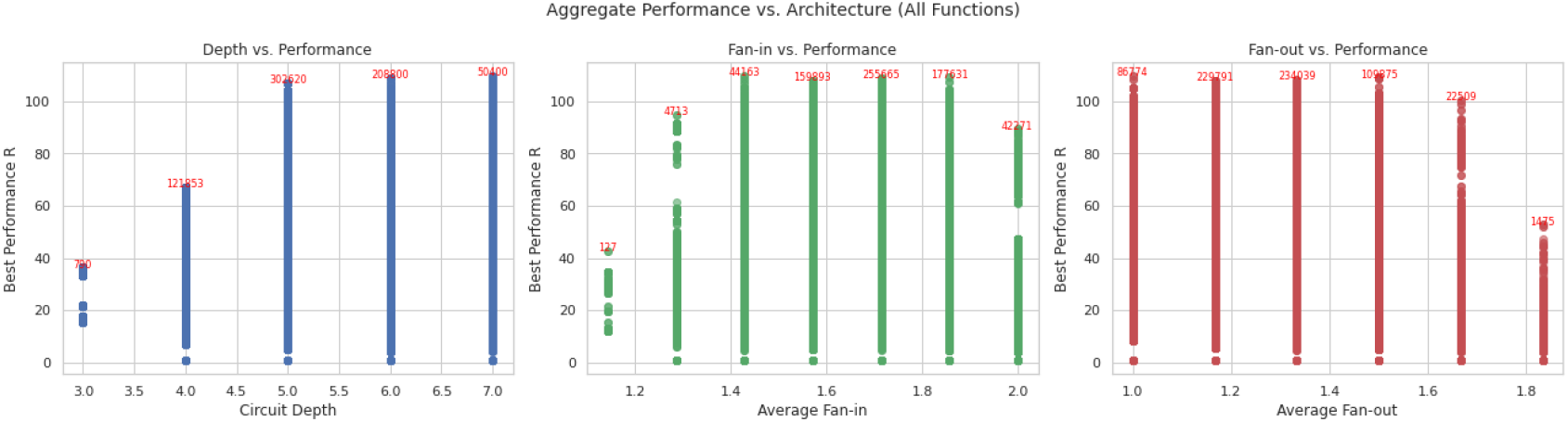
B. Aggregate plots comparing *µ* to circuit depth, average fan-in, and average fan-out for all 2-input Boolean functions using *k* = 7 gates

**Figure 22:** Over all 2-input Boolean functions using *k* = 7 gates, we observe that performance generally increases as average fan-in increases and average fan-out decreases. A higher depth allows for a higher average fan-in resulting in larger *µ* with a deeper circuit.

## References

[1] H. H. McAdams and L. Shapiro, “Circuit simulation of genetic networks,” Science, vol. 269, no. 5224, pp. 650–656, 1995.

[2] A. S. Khalil and J. J. Collins, “Synthetic biology: applications come of age,” Nat. Rev. Genet., vol. 11, no. 5, pp. 367–379, 2010.

[3] J. H. Esensten, J. A. Bluestone, and W. A. Lim, “Engineering therapeutic T cells: from synthetic biology to clinical trials,” Annu. Rev. Pathol.: Mech. Dis., vol. 12, pp. 305–330, 2017.

[4] P.-F. Xia, H. Ling, J. L. Foo, and M. W. Chang, “Synthetic genetic circuits for programmable biological functionalities,” Biotechnol. Adv., 2019.

[5] S. Itzkovitz, T. Tlusty, and U. Alon, “Coding limits on the number of transcription factors,” BMC genomics, vol. 7, no. 1, p. 239, 2006.

[6] A. Garg, J. J. Lohmueller, P. A. Silver, and T. Z. Armel, “Engineering synthetic TAL effectors with orthogonal target sites,” Nucleic Acids Res., vol. 40, no. 15, pp. 7584–7595, 2012.

[7] B. C. Stanton, A. A. Nielsen, A. Tamsir, K. Clancy, T. Peterson, and C. A. Voigt, “Genomic mining of prokaryotic repressors for orthogonal logic gates,” Nat. Chem. Biol., vol. 10, no. 2, p. 99, 2014.

[8] L. S. Qi, M. H. Larson, L. A. Gilbert, J. A. Doudna, J. S. Weissman, A. P. Arkin, and W. A. Lim, “Repurposing CRISPR as an RNA-guided platform for sequence-specific control of gene expression,” Cell, vol. 152, no. 5, pp. 1173–1183, 2013.

[9] L. A. Gilbert, M. H. Larson, L. Morsut, Z. Liu, G. A. Brar, S. E. Torres, N. Stern-Ginossar, O. Brandman, E. H. Whitehead, J. A. Doudna, et al., “CRISPR-mediated modular RNA-guided regulation of transcription in eukaryotes,” Cell, vol. 154, no. 2, pp. 442–451, 2013.

[10] A. A. Nielsen and C. A. Voigt, “Multi-input CRISPR/Cas genetic circuits that interface host regulatory networks,” Mol. Syst. Biol., vol. 10, no. 11, 2014.

[11] M. W. Gander, J. D. Vrana, W. E. Voje, J. M. Carothers, and E. Klavins, “Digital logic circuits in yeast with CRISPR-dCas9 NOR gates,” Nat. Commun., vol. 8, no. 1, pp. 1–11, 2017.

[12] J. M. Rock, F. F. Hopkins, A. Chavez, M. Diallo, M. R. Chase, E. R. Gerrick, J. R. Pritchard, G. M. Church, E. J. Rubin, C. M. Sassetti, et al., “Programmable transcriptional repression in mycobacteria using an orthogonal CRISPR interference platform,” Nat. Microbiol., vol. 2, no. 4, pp. 1–9, 2017.

[13] L. Cui, A. Vigouroux, F. Rousset, H. Varet, V. Khanna, and D. Bikard, “A CRISPRi screen in E. coli reveals sequence-specific toxicity of dCas9,” Nat. Commun., vol. 9, no. 1, pp. 1–10, 2018.

[14] S. Zhang and C. A. Voigt, “Engineered dCas9 with reduced toxicity in bacteria: implications for genetic circuit design,” Nucleic Acids Res., vol. 46, no. 20, pp. 11115–11125, 2018.

[15] S. Cho, D. Choe, E. Lee, S. C. Kim, B. Palsson, and B.-K. Cho, “High-level dCas9 expression induces abnormal cell morphology in Escherichia coli,” ACS Synth. Biol., vol. 7, no. 4, pp. 1085–1094, 2018.

[16] B. M. Markus, G. W. Bell, H. A. Lorenzi, and S. Lourido, “Optimizing systems for Cas9 expression in Toxoplasma gondii,” mSphere, vol. 4, no. 3, pp. e00386–19, 2019.

[17] A. Tamsir, J. J. Tabor, and C. A. Voigt, “Robust multicellular computing using genetically encoded NOR gates and chemical ‘wires’,” Nature, vol. 469, no. 7329, pp. 212–215, 2011.

[18] D. Ausländer, S. Ausländer, X. Pierrat, L. Hellmann, L. Rachid, and M. Fussenegger, “Programmable full-adder computations in communicating three-dimensional cell cultures,” Nat. Methods, vol. 15, no. 1, p. 57, 2018.

[19] M. A. Al-Radhawi, A. P. Tran, E. A. Ernst, T. Chen, C. A. Voigt, and E. D. Sontag, “Distributed implementation of boolean functions by transcriptional synthetic circuits,” ACS Synthetic Biology, vol. 9, no. 8, pp. 2172–2187, 2020.

[20] P.-Y. Chen, Y. Qian, and D. Del Vecchio, “A model for resource competition in crispr-mediated gene repression,” in 2018 IEEE Conference on Decision and Control (CDC), pp. 4333–4338, IEEE, 2018.

[21] H.-H. Huang, M. Bellato, Y. Qian, P. Cárdenas, L. Pasotti, P. Magni, and D. Del Vecchio, “dcas9 regulator to neutralize competition in crispri circuits,” Nature communications, vol. 12, no. 1, pp. 1–7, 2021.

[22] J. Santos-Moreno, E. Tasiudi, J. Stelling, and Y. Schaerli, “Multistable and dynamic crispri-based synthetic circuits,” Nat. Commun., vol. 11, no. 1, pp. 1–8, 2020.

[23] A. A. Nielsen, B. S. Der, J. Shin, P. Vaidyanathan, V. Paralanov, E. A. Strychalski, D. Ross, D. Densmore, and C. A. Voigt, “Genetic circuit design automation,” Science, vol. 352, no. 6281, p. aac7341, 2016.

[24] P. Érdi and J. Tóth, Mathematical models of chemical reactions: theory and applications of deterministic and stochastic models. Manchester University Press, 1989.

[25] S. Clamons and R. Murray, “Modeling predicts that crispr-based activators, unlike crispr-based repressors, scale well with increasing grna competition and dcas9 bottlenecking,” bioRxiv, p. 719278, 2019.

[26] J. A. Brophy and C. A. Voigt, “Principles of genetic circuit design,” Nature methods, vol. 11, no. 5, pp. 508–520, 2014.

[27] P. Ellison, M. Feinberg, H. Ji, and D. Knight, “Chemical reaction network toolbox,” Available online a t http://www.crnt.osu.edu/CRNTWin, 2011.

